# Gut bacteria induce heterologous immune priming in *Rhodnius prolixus* encompassing both humoral and cellular immune responses

**DOI:** 10.1101/2025.01.31.635857

**Authors:** Carissa A. Gilliland, Kevin J. Vogel

## Abstract

Insects lack the adaptive, antibody mediated responses of vertebrates, yet they possess a robust innate immune system capable of defending the host against pathogens. Immune priming has been observed in multiple insect species, wherein exposure to a pathogen provides protection against subsequent infections by the pathogen. Less frequently, heterologous immune priming has been observed where exposure to one bacterial species provides protection against other species. We determined that *Rhodococcus rhodnii*, a gut symbiont of the kissing bug *Rhodnius prolixus,* induces a strong heterologous immune priming effect, while axenic bugs lacking any gut bacteria are highly susceptible to pathogens in their hemolymph. Commensal *Escherichia coli* provides a less robust protective effect than *R. rhodnii*. *R. rhodnii* must be alive within the insect as dead bacteria do not stimulate immune priming and pathogen resistance. Removal of *R. rhodnii* from the gut reduces resistance to pathogens while restoring it to otherwise axenic bugs improves resistance to pathogens, though not completely. *R. rhodnii* and *E. coli* activate both the Imd and Toll pathways, indicating cross-activation of the pathways and demonstrating the canonical *Drosophila* immune response has diverged in Hemiptera. Silencing of either pathway leads to a loss of the protective effect. Several antimicrobial peptides are induced in the fat body by presence of gut bacteria. When *E. coli* is in the gut, expression of antimicrobial peptides is often higher than when *R. rhodnii*, though *R. rhodnii* stimulates proliferation of hemocytes and induce a stronger melanization response. Hemolymph from *R. rhodnii* bugs has a greater ability to convert the melanin precursor DOPA to melanization products than axenic or *E. coli*-harboring bugs. These results demonstrate that *R. rhodnii’s* benefits to its host extend beyond nutritional provisioning, playing an important role in the host immune system.

**Author Summary:** Insects often form beneficial relationships with bacteria allowing them to eat nutritionally deficient diets. In insects that only consume blood, symbionts are necessary to provide B vitamins absent in the host diet. There is a growing appreciation that in some of these symbiotic associations, the bacteria provide services beyond nutrition. We show that in kissing bugs, which feed exclusively on vertebrate blood and require bacterial symbionts for development, these symbiotic bacteria are important in activating the insect immune system. Insects with no gut bacteria are highly susceptible to infection and cannot mount an effective immune response. The bacteria reside exclusively in the insect gut yet protect against infections in the rest of the insect’s body. The bacteria must be alive to prime the immune system, and the response is dependent on the species of bacteria in the gut, with symbiotic bacteria providing stronger protection against infection and inducing a broader array of immune responses than commensal bacteria. This study expands our understanding of the role of beneficial bacteria in insect immunity and demonstrates that immune systems differ between major groups of insects.

## Introduction

Though they lack the adaptive, acquired immunity seen in vertebrates, insects possess a robust and well-developed innate immune system [1] allowing them to mount a strong and specific defense against pathogens [2–6]. An immune response is initiated when an insect pattern recognition receptor recognizes a conserved motif or microbially-associated molecular pattern (MAMP) that is present in microbes but not found in insects [7]. Pathogen recognition triggers the activation of signal transduction pathways that amplify immune responses and activates immune cascades corresponding to the specific pathogen [8]. Immune effector mechanisms include production of antimicrobial peptides, phagocytosis, melanization, encapsulation, lysis, and others [4,8–10].

Insect immunity is often characterized into two response systems, humoral and cellular, though this division is for the ease of discussion as these systems are largely connected. Humoral defenses consist mainly of soluble effector molecules mainly derived from the fat body, an organ analogous to the vertebrate liver with both nutritional and immunological functions [11–14]. The humoral immune cascade can be activated via three different pathways: the Toll pathway, which recognizes Gram-positive bacteria and fungi, the immune deficiency (Imd) pathway, which recognizes Gram-negative bacteria, and the Jak/Stat pathway, which recognizes viruses [4]. Cellular defenses are mediated by hemocytes circulating in the hemolymph [10]. These cells can sequester and kill microbes via phagocytosis, encapsulation, or deposition of toxic melanin (melanization).

The insect gut is considered a crucial organ in insect immune defense as it is exposed to environmental microbes, including pathogens, through feeding [15]. When an insect ingests a pathogen, the immune pathways may be activated and produce different AMPs to maintain gut homeostasis [16]. In addition to encountering pathogenic bacteria, insects often maintain beneficial associations with microbes they rely upon for successful development and reproduction. Some symbiotic bacteria have been shown to benefit their host through enhancing the immune response to pathogens. Tsetse flies harbor intracellular *Wigglesworthia* in bacteriocytes which provide essential B vitamins to the host. Flies cleared of *Wigglesworthia* are more likely to succumb to infection with normally non-pathogenic *Escherichia coli* compared to their symbiotic counterparts [17]. In the bean bug, *Riptortus pedestris,* their symbiont *Caballeronia* and other commensal bacteria increase host immune competence when challenged with an entomopathogen [18,19]. The microbiota of honeybees, *Apis mellifera*, have also been shown to be an essential component of the host immune response to pathogens [20].

The role of bacterial symbionts in the immune responses of kissing bugs (Hemiptera: Reduviidae) has not been well studied. Like all other obligately and exclusively hematophagous arthropods, kissing bugs rely on microbes to provide essential B vitamins that are deficient in vertebrate blood, and these microbes are essential for host development [21,22]. The kissing bug, *Rhodnius prolixus,* was discovered to house a free-living, Gram-positive bacterium in the lumen of the gut, *Rhodococcus rhodnii* [23–25]. This bacterium was demonstrated to be beneficial to the host as its removal via surface sterilization of eggs results in increased mortality, increased development time, and failure to reach adulthood [22,25,26].

Surveys utilizing 16S rDNA amplicon sequencing of kissing bug gut microbiomes highlight that other microbes are present in wild populations and that *Rhodococcus* is not ubiquitous across species, especially in the genus *Triatoma* [27–37]. These studies have shown that kissing bug microbiomes comprise a limited diversity of bacteria that can range from dozens to hundreds of members. Although kissing bugs can house multiple species of bacteria it has been demonstrated that *R. rhodnii* is necessary and sufficient to support development and reproduction in *R. prolixus* [27]. When specifically looking at members of the genus *Rhodnius*, others have found that *R. rhodnii* is ubiquitous among the genus, suggesting it is an important co-evolved symbiont of this group [38]. In addition to *R. rhodnii*, other bacteria have been found in guts of *Rhodnius* sp. [39–41] though their specific roles are not known. The diversity of microbes within and amongst these studies suggests that the *Rhodnius* microbiome contains at least some commensal organisms. Our previous work indicated that non-*Rhodococcus* species including *Escherichia coli* could at least temporarily colonize the gut of *R. prolixus*, and that some proportion of these insects were able to successfully reach adulthood and reproduce, though not as many as insects harboring *R. rhodnii* [1]. Insects harboring exclusively *E. coli* did not exhibit signs of infection, and *E. coli* remained at a much lower titer in the gut than *R. rhodnii*. These results suggested that in the gut of *R. prolixus*, *E. coli* could exist as a non-pathogenic resident.

The observation that *R. prolixus* can harbor multiple species of bacteria in its gut led us to wonder how the host immune system is impacted by the presence of different microbes in the gut. Using our ability to produce axenic insects (germ free) and those with a defined microbiota (gnotobiotic), we compared the host’s immune response to pathogens in these two states and explored the mechanisms, if any, underlying resistance to pathogens. Our results highlight how symbionts and commensals can play distinct roles in shaping the host immune response.

## Results

### The Presence of Bacteria Protects *R. prolixus* Against Bacterial Infection

To investigate the effects of gut bacteria on host immune function, we removed all bacteria through surface sterilization of eggs using our previously described protocol [28]. Bugs were either reared in axenic conditions (Rpro^Axn^), or experimentally infected through a blood meal with *E. coli* MG1655 (Rpro^Ec^), or their symbiont *R. rhodnii* ATCC 35071 (Rpro^Rr^). These bugs were injected with 10^6^ CFUs of *E. coli*, *R. rhodnii*, the Gram-positive *Micrococcus luteus* or sterile saline into the hemolymph of unfed 4^th^ instar nymphs and survival after infection was monitored. Almost all Rpro^Axn^ individuals died within 5 days after injection with either *E. coli* (9% survival, Fig. 1A) or *M. luteus* (17% survival, Fig. 1A). In contrast, Rpro^Rr^ bugs showed significantly higher survival after *E. coli* or *M. luteus* infection (79% and 68% survival, respectively) than Rpro^Axn^ (p < 0.0001 for both pathogens, log-rank test). Rpro^Ec^ bugs had significantly higher survival rates compared to Rpro^Axn^ individuals for both challenges (52% survival, p < 0.0001, and 50% survival, p = 0.003, respectively, log-rank test, Fig. 1A) and lower survival relative to Rpro^Rr^ challenged with *E. coli* (p = 0.018, log-rank test) but not when challenged with *M. luteus* (p = 0.07, log-rank test). To test if the mortality we saw was due to bacterial infection or due to wounding alone, we also injected Rpro^Axn^, Rpro^Ec^, and Rpro^Rr^ insects with sterile saline. All insects subject to sterile saline injection had high survival and were not significantly different from each other (p > 0.05, log-rank test, Fig. 1A).

**Figure 1:**
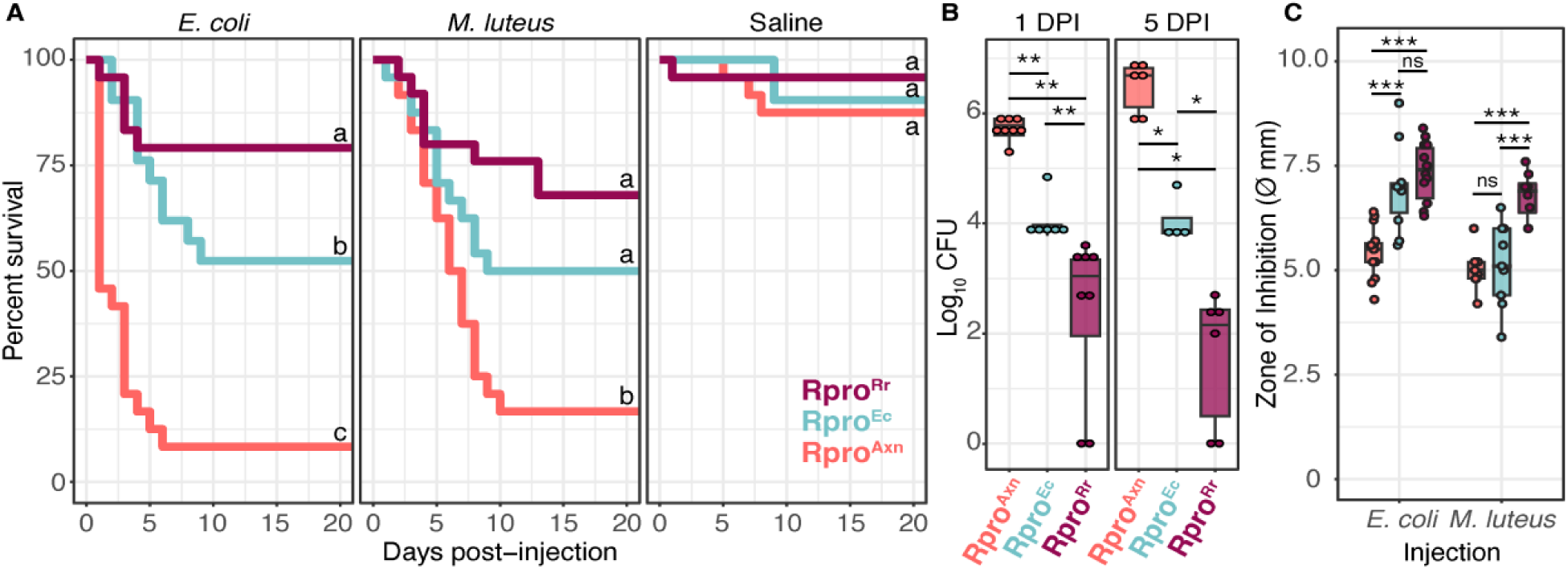
Gut bacteria promote immune priming against bacterial infection in *R. prolixus*. **(A)** Survival curves of Rpro^Axn^, Rpro^Ec^, and Rpro^Rr^ after injection with 10^6^ CFU of *E. coli, M. luteus,* or sterile saline into their hemocoel. Rpro^Rr^ and Rpro^Ec^ had significantly higher survival than Rpro^Axn^ regardless of the bacteria used for challenge (p < 0.0001, log-rank test), while there was no difference in survival of bugs injected with sterile saline (p ≥ 0.15, log-rank test) indicating that the presence of gut microbes plays a critical role in insect defense against pathogens. When challenged with *E. coli* there was a significant difference in survival (26.8%) between Rpro^Rr^ and Rpro^Ec^ (p = 0.018, log-rank test). The difference in survival between Rpro^Rr^ and Rpro^Ec^ was smaller (18%) and approached but did not reach significance when challenged with *M. luteus* (p = 0.072, log-rank test). Lines connected by different letters are significantly different (p < 0.05, log-rank test). **(B)** Gnotobiotic *R. prolixus* limit the growth of *E. coli* in the hemocoel. Boxplots *of E. coli* CFUs in *R. prolixus* hemolymph collected at 1 and 5 DPI. Points represent individual bugs. Rpro^Rr^ bugs had fewer *E. coli* CFU at both 1 and 5 DPI than Rpro^Ec^ or Rpro^Axn^. Rpro^Ec^ had fewer *E. coli* CFU at both 1 and 5 DPI than Rpro^Axn^ (** p < 0.002, * p < 0.05, Wilcoxon test). **(C)** Hemolymph from Rpro^Rr^ bugs suppresses growth of both *E. coli* and *M. luteus in vitro* more than Rpro^Ec^ or Rpro^Axn^ bugs. ***p < 0.001, Tukey’s HSD.

Our survival curves indicate that bugs reared with bacteria in their gut can survive infection with *E. coli* or *M. luteus* while nearly all Rpro^Axn^ die after infection. Insects can overcome infection through either resistance - killing or clearing of a pathogen – or through tolerance – minimizing fitness costs of infection without reducing the pathogen load. To determine if the observed difference in survival was due to tolerance or resistance, we measured *E. coli* titer in the hemolymph after infection over time. Because both the *E. coli* and *M. luteus* treatment groups had similar mortality rates we chose to further investigate only *E. coli* for this experiment. We injected bugs with 10^6^ CFU of live, kanamycin-resistant *E. coli,* then collected hemolymph at 1-and 5-days post injection. Hemolymph was spread onto plates containing kanamycin to calculate CFUs of the kanamycin-resistant *E. coli* present in the hemolymph. The presence of bacteria in the gut had a significant impact on the number of Kan^R^ *E. coli* hemolymph (F_2,32_ = 123.2, p < 0.0001, aligned-rank transformed ANOVA). One day post injection with *E. coli*, Rpro^Rr^ insects had on average 1.6 x 10^3^ CFUs of *E. coli*, Rpro^Ec^ had 1.82 x 10^4^, and Rpro^Axn^ individuals had 4.6 x 10^5^ CFUs, higher than either gnotobiotic treatment (p = 0.002, Wilcoxon signed-rank test, Fig. 1B). Five days after injection with *E. coli* most Rpro^Axn^ had died; those who remained alive had on average 4.3 x 10^6^ CFUs of *E. coli.* Rpro^Ec^ bugs had significantly fewer *E. coli* than Rpro^Axn^ bugs (1.8 x 10^4^ CFUs, p = 0.014, Wilcoxon signed-rank test, Fig. 1B), while Rpro^Rr^ bugs had the least *E. coli* in their hemolymph of any treatment (2 x 10^2^ CFUs, p = 0.014, Wilcoxon signed-rank test). The identity of the gut bacteria influences the extent to which the gnotobiotic bugs can clear the bacteria from their hemolymph, with Rpro^Rr^ bugs having significantly fewer bacteria at day 5 than Rpro^Ec^ bugs (p = 0.014, Wilcoxon signed-rank test). Though neither gnotobiotic treatment completely cleared *E. coli* from their hemolymph during the observation window, they appear to suppress the pathogen to a level that is survivable.

We next sought to determine if the decrease in bacteria in the hemocoel of challenged Rpro^Axn^, Rpro^Ec^, and Rpro^Rr^ bugs was due to bacterial killing via antimicrobial factors in the hemolymph. To explore if presence of gut bacteria results in differences in antimicrobial activity in the bugs’ hemolymph, we conducted a zone of inhibition assay [42,43]. We injected Rpro^Axn^, Rpro^Ec^, and Rpro^Rr^ 4^th^ instar bugs with 10^6^ cells of either heat-killed *E. coli* or *M. luteus.* We used heat-killed bacteria for this assay to avoid bacterial growth on the plates that could potentially confound our zone of inhibition measurements. Rpro^Rr^ and Rpro^Ec^ bugs had significantly larger zones of inhibition when infected with *E. coli* than Rpro^Axn^ bugs (p < 0.0001, Tukey’s HSD, Fig. 1C). Interestingly, only Rpro^Rr^ bugs had significantly larger zones of inhibition after being infected with *M. luteus* (p < 0.0001, Tukey’s HSD, Fig. 1C). These results indicate that Rpro^Rr^ and Rpro^Ec^ bugs have higher antimicrobial activity in their hemolymph when challenged with Gram-negative *E. coli* while only Rpro^Rr^ bugs had higher antimicrobial activity after infection with the Gram-positive *M. luteus,* possibly due to activation of the Toll pathway in Rpro^Rr^. Naïve individuals did not show a significant difference in antimicrobial activity from each other, suggesting this is an induced response (Fig. S1). These data demonstrate the presence of gut bacteria promotes resistance to pathogens in *R. prolixus* through a reduction in the number of bacteria in the hemolymph of gnotobiotic bugs but not Rpro^Axn^ bugs.

### The Protective Effect of *R. rhodnii* Requires Live Bacteria and is Lost Upon Removal

Insects sense bacteria through detection of MAMPs such as peptidoglycan, lipopolysaccharides, and lipoteichoic acid, which activate signaling cascades via serine proteases. We therefore asked whether the immune priming effect of *R. rhodnii* was primarily due to the presence of MAMPs, and so we fed Rpro^Axn^ bugs blood meals that were spiked with an amount of heat-killed *R. rhodnii* that was equivalent to 10^8^ CFUs/bug (Rpro^Axn+HK Rr^) from the first blood meal until they developed into 4^th^ instars, then challenged them as before with injection of *E. coli*. All Rpro^Axn+HK Rr^ bugs died within the 10-day window and their survival was not statistically different from the survival of Rpro^Axn^ bugs (p = 1.00, Cox proportional hazards model, Fig. 2A). From these results we conclude that live bacteria are necessary for immune priming in this system.

**Figure 2:**
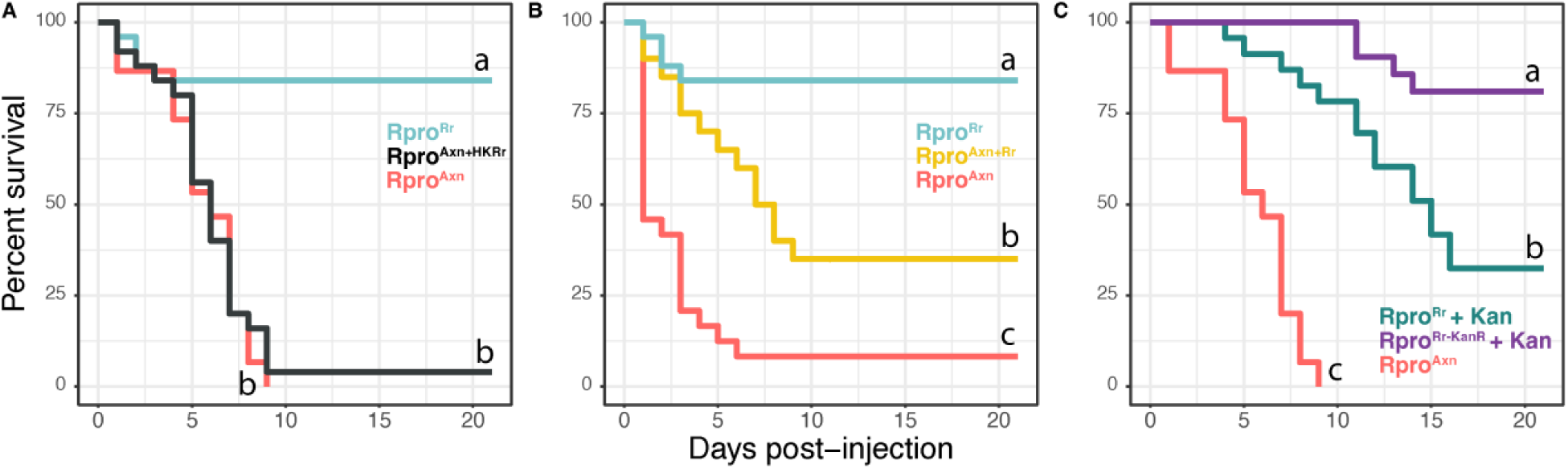
*R. prolixus* requires live bacteria to mount an effective immune response. Within a panel, lines not connected by the same letter are significantly different (p < 0.001, log-rank test). **(A)** Survival after bacterial injection of 4^th^ instar Rpro^Axn^ nymphs fed heat-killed *R. rhodnii* throughout development (Rpro^Axn + HK Rr^) was similar to 4^th^ instar Rpro^Axn^ nymphs not fed bacteria (p = 0.69, log-rank test). Survival of Rpro^Rr^ was significantly higher (p < 0.0001, log-rank test). **(B)** Survival after immune challenge of Rpro^Axn^ bugs after restoration of *R. rhodnii* (Rpro^Axn+ Rr^). Restoring *R. rhodnii* via a blood meal significantly increases survival relative to axenic nymphs (p < 0.0001, log-rank test) but there was still significantly lower survival in the Rpro^Axn + Rr^ than in Rpro^Rr^ (p < 0.0001, log-rank test). **(C)** Clearance of *R. rhodnii* from Rpro^Rr^ significantly reduced survival following immune challenge with *E. coli*. Rpro^Rr-KanR^ bugs treated with kanamycin retained their immune priming effect. Different letters represent significant differences between treatments (p < 0.001, log-rank test).

The main function of *R. rhodnii* is thought to be supplementation of B vitamins to the host [22,44]. These nutrients are important in numerous essential processes in the host and their absence throughout Rpro^Axn^ development may underlie the higher survival of Rpro^Rr^ following immune challenge. Alternatively, the protective effect of *R. rhodnii* may be independent of its nutrient provisioning services. If the effect were nutritional, we would expect that removal of *R. rhodnii* from Rpro^Rr^ bugs shortly before an immune challenge would not have a dramatic effect on survival, while if the effect were primarily immune priming, addition of *R. rhodnii* to Rpro^Axn^ bugs shortly before challenge would lead to increased survival relative to Rpro^Axn^ individuals.

To investigate this, we fed 3^rd^ instar Rpro^Axn^ nymphs a blood meal containing 10^6^ CFU/mL of *R. rhodnii*. Nymphs developed into 4^th^ instars and a subset were sacrificed and qPCR was used to confirm that sacrificed nymphs harbored *R. rhodnii* (Fig. S2). Two weeks after molting, insects were injected with live *E. coli* as previously outlined. Rpro^Axn+Rr^ bugs had higher survival than Rpro^Axn^ bugs (Fig. 2B, p = 0.0047, Cox proportional hazards) but still suffered increased mortality compared to Rpro^Rr^ individuals. Interestingly, when we removed *R. rhodnii* from Rpro^Rr^ bugs by feeding kanamycin to 3^rd^ instar insects via the blood meal (Rpro^Rr^ + Kan), they also suffered from increased mortality compared to bugs that were fed kanamycin but had been inoculated with kanamycin resistant *R. rhodnii* (Rpro^Rr-KanR^ + Kan, Fig. 2C). Thus, we conclude that the nutritional role of *R. rhodnii* in *R. prolixus* is not sufficient to explain the observed immune priming effects, but that nutrient provisioning may contribute to the protective effects of *R. rhodnii*. We did not test this in Rpro^Ec^ bugs, as their B vitamin synthesis capabilities are similar to *R. rhodnii*, and we do not anticipate that nutritional provisioning by *E. coli* would be able to rescue immune priming when *R. rhodnii* could not.

### Immune Priming by Gut Bacteria is Dependent on Both Toll and Imd Pathways

Immune responses to bacteria in insects are thought to be mediated by two major pathways, Toll and Imd [44]. Our zone of inhibition assays suggested that some factor(s) in the hemolymph are involved in suppressing bacterial proliferation, which may be regulated by Toll or Imd. To investigate whether the Toll or Imd pathways are essential to protecting kissing bugs against bacterial infection, we first measured expression of two transcription factors, *dorsal* and *relish*, which drive expression of Toll and Imd response genes including antimicrobial peptides (AMPs). For both genes, we assessed expression via qPCR in the fat body of naïve insects and insects that were injected with live *E. coli* or *M. luteus* as described above. We tested Rpro^Rr^, Rpro^Ec^, and Rpro^Axn^ bugs, and found that naïve, Rpro^Axn^ bugs had low expression of both dorsal and relish (Fig. 3A, B).

**Figure 3:**
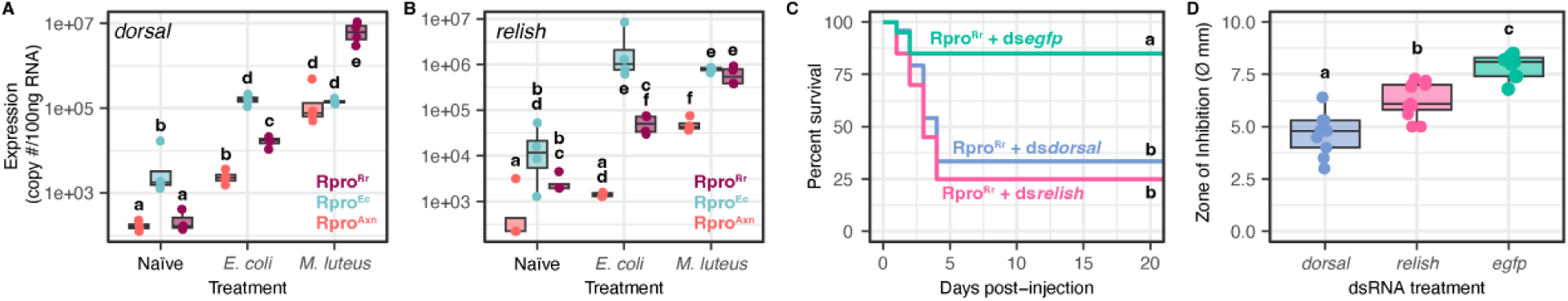
Immune priming by gut bacteria is dependent on Toll and Imd pathways. **(A)** Expression of the Toll pathway transcription factor *dorsal* in Rpro^Rr^, Rpro^Ec^, and Rpro^Axn^ in naïve bugs or bugs challenged with *E. coli* or *M. luteus* injection. Bars connected by different letters are significantly different (p < 0.05, Tukey’s HSD). **(B)** Expression of the Imd pathway transcription factor *relish* in Rpro^Rr^, Rpro^Ec^, and Rpro^Axn^ in naïve bugs or bugs challenged with *E. coli* or *M. luteus* injection. Bars connected by different letters are significantly different (p < 0.05, Tukey’s HSD). **(C)** Survival curves of Rpro^Rr^ bugs treated with dsRNA against *dorsal, relish*, or a control *egfp* sequence demonstrate that silencing of *relish* or *dorsal* via RNAi significantly reduced the survival of Rpro^Rr^ bugs, indicating that these pathways are necessary for *R. rhodnii*-mediated immune priming. Different letters indicate treatments with significantly different survival (p < 0.0001, log-rank test). **(D)** Silencing *dorsal* or *relish* reduces the bacteriostatic factors in hemolymph of Rpro^Rr^. Different letters indicate treatments with significantly different survival (p < 0.0001, Tukey’s HSD).

For each gene, there was a highly significant effect of the gnotobiotic state of the bugs, the immune challenge, and the interaction between state and challenge (see Table S2 for statistical details). Naïve Rpro^Ec^ bugs had higher expression of *dorsal* and *relish* than Rpro^Axn^, indicating that *E. coli* in the gut stimulates expression of both transcription factors independent of immune challenge via injection. Naïve Rpro^Rr^ bugs had low expression of *dorsal*, equivalent to Rpro^Axn^ bugs but had higher expression of *relish* than Rpro^Axn^. In bugs challenged with injection of *E. coli*, both *dorsal* and *relish* expression were elevated in their respective gnotobiotic bugs relative to Rpro^Axn^. *M. luteus* also stimulated robust expression of *relish* and *dorsal*, with Rpro^Rr^ bugs having the strongest expression of *dorsal* while *relish* expression was high in both Rpro^Rr^ and Rpro^Ec^.

From our expression analysis of *dorsal* and *relish*, it is clear that there is significant cross-activation of the Toll and Imd pathways, with both Gram-negative and Gram-positive bacteria activating each. The Toll pathway, and by extension *dorsal*, is thought to primarily respond to Gram-positive bacteria while Imd and *relish* are thought to be activated by Gram-negative bacteria, but significant crosstalk between these pathways has been observed in other hemipterans [45]. Absence of gut bacteria does not eliminate the ability of bugs to mount an immune response, but gnotobiotic bugs almost always have higher expression of these genes. In all bugs tested, both *dorsal* and *relish* can be induced by the presence of bacteria in the hemolymph, but we were surprised to see that Rpro^Ec^ often induces higher expression of *dorsal* and *relish* than Rpro^Rr^, given the latter’s stronger protective effect.

We next asked whether the immune priming effect of gut bacteria is dependent on the Toll or Imd pathway. We suppressed each pathway via RNAi knockdown of the transcription factors *relish* (Imd) or *dorsal* (Toll). RNAi of *relish* or *dorsal* was confirmed via qPCR revealing over a 90% reduction in *dorsal* or *relish* transcripts after injection with dsRNA (Fig. S3). After injection with dsRNA, 10^6^ CFUs of *E. coli* were injected into the bugs hemocoel, and survival rates were measured (Fig. 3C). Rpro^Rr^ treated with either *dorsal* or *relish* dsRNA succumbed to *E. coli* infection at a higher rate than Rpro^Rr^ individuals treated with ds*egfp* (25% and 34% survival respectively, p = 0.002, p = 0.0004, respectively, log-rank test).

We then examined the change in antimicrobial activity in *relish* or *dorsal*-silenced *R. prolixus* by comparing the inhibitory effects of hemolymph on microbial growth using the zone of inhibition assay described above, though we only tested the effect of hemolymph from dsRNA-treated bugs against *E. coli*. There was a significant effect of silencing either *relish* or *dorsal* as silenced bugs had smaller zones of inhibitions compared to the control ds*egfp*-injected group (Fig. 3D, p < 0.0007, p < 0.0001, Tukey’s HSD). The decrease in survival after challenge with *E. coli* in either *relish* or *dorsal* knockdown bugs suggests immune pathway crosstalk in *R. prolixus*, as silencing of the Toll pathway transcription factor *dorsal* led to significantly higher mortality following challenge with Gram-negative bacteria. The reduction in antimicrobial activity in hemolymph following knockdown of *dorsal* or *relish* suggests that the activation of the Toll and Imd pathways by *R. rhodnii* has functional consequences for host immune responses.

### Expression of Immune-Related Genes is Influenced by the Gut Microbiome

Our experiments demonstrated that *dorsal* and *relish* were important mediators of the immune priming effect seen in Rpro^Rr^. We next wanted to see if antimicrobial peptides were upregulated in gnotobiotic insects. To test this, we evaluated the fat body expression profiles of AMPs in 4^th^ instar Rpro^Axn^, Rpro^Ec^, and Rpro^Rr^ bugs that were either uninfected or 1-day post-inoculation with either *E. coli* or *M. luteus*. We measured the expression of the AMPs *prolixicin*, two defensins (RPRC012182 and RPRC012184, subsequently referred to as *defensin82*, and *defensin84*), and a lysozyme (RPRC015442 subsequently referred to as *lysozyme 42*, Fig. 4).

**Figure 4:**
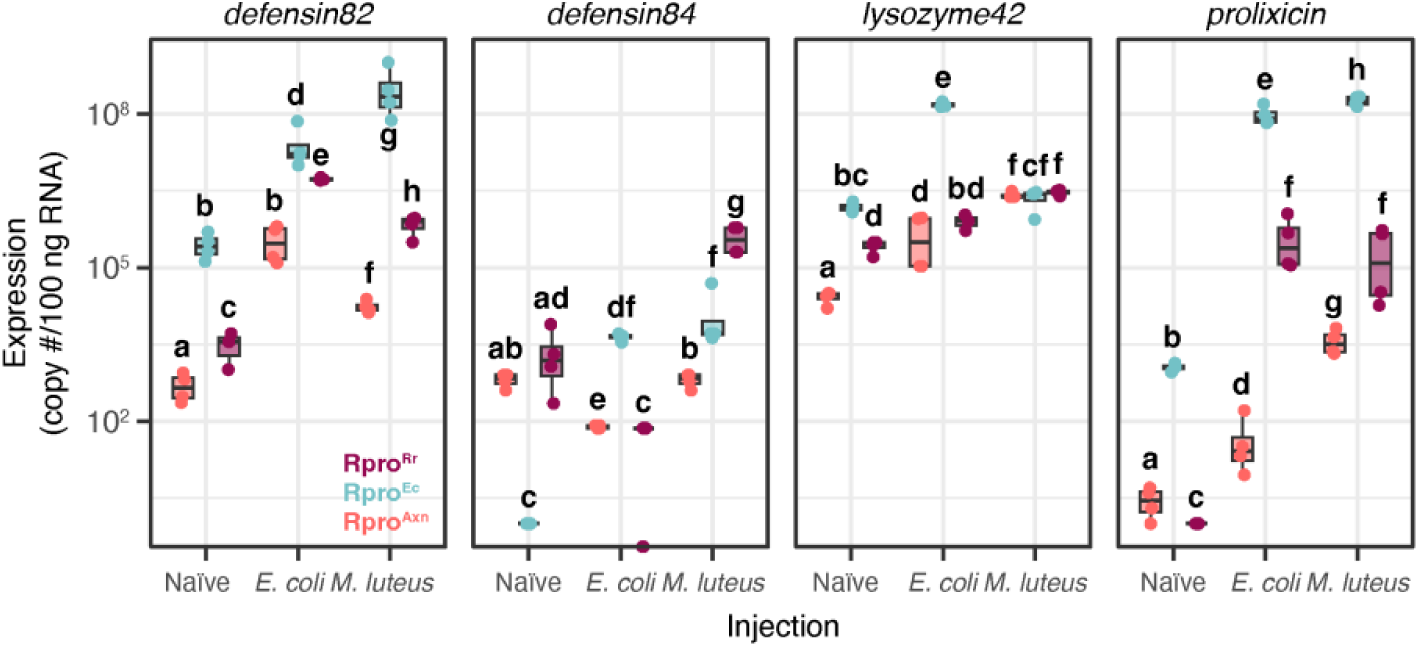
Antimicrobial peptide (AMP) genes are differentially expressed among naïve, Rpro^Ec^, and Rpro^Rr^ bugs. Within each AMP panel, bars connected by different letters are significantly different (p < 0.05, Tukey’s HSD).

The interaction of gnotobiotic state x injection was highly significant for all genes tested (p < 0.0001, aligned ranks transformation ANOVA for non-parametric interactions).

Our expression analysis of immune genes revealed several interesting patterns. First, Rpro^Axn^ insects often had lower expression of AMP genes regardless of the immune challenge (Fig. 5), which may be a consequence of lower *relish* and *dorsal* expression in Rpro^Axn^ bugs (Fig. 3A, B). The overall reduced expression of immune genes may partially explain the heightened susceptibility of Rpro^Axn^ to pathogens. A second, surprising pattern is that Rpro^Ec^ bugs often have higher expression of AMPs than Rpro^Rr^ bugs, despite Rpro^Rr^ having a significantly higher survival rate when challenged with pathogens. In the Rpro^Ec^ bugs challenged with *E. coli*, expression of all immune genes except *defensin84* were significantly higher than when Rpro^Rr^ bugs were challenged with *E. coli*, indicative of a strong immune priming effect of *E. coli* in the gut against *E. coli* in the hemolymph (Fig. 4, p < 0.05, Tukey’s HSD). Rpro^Rr^ bugs exhibited strong induction of immune gene expression in response to bacterial challenge relative to naïve Rpro^Rr^ or Rpro^Axn^, apart from *defensin84*, suggesting that humoral immune responses are important in the heterologous immune priming seen in Rpro^Rr^ following pathogen challenge.

**Figure 5:**
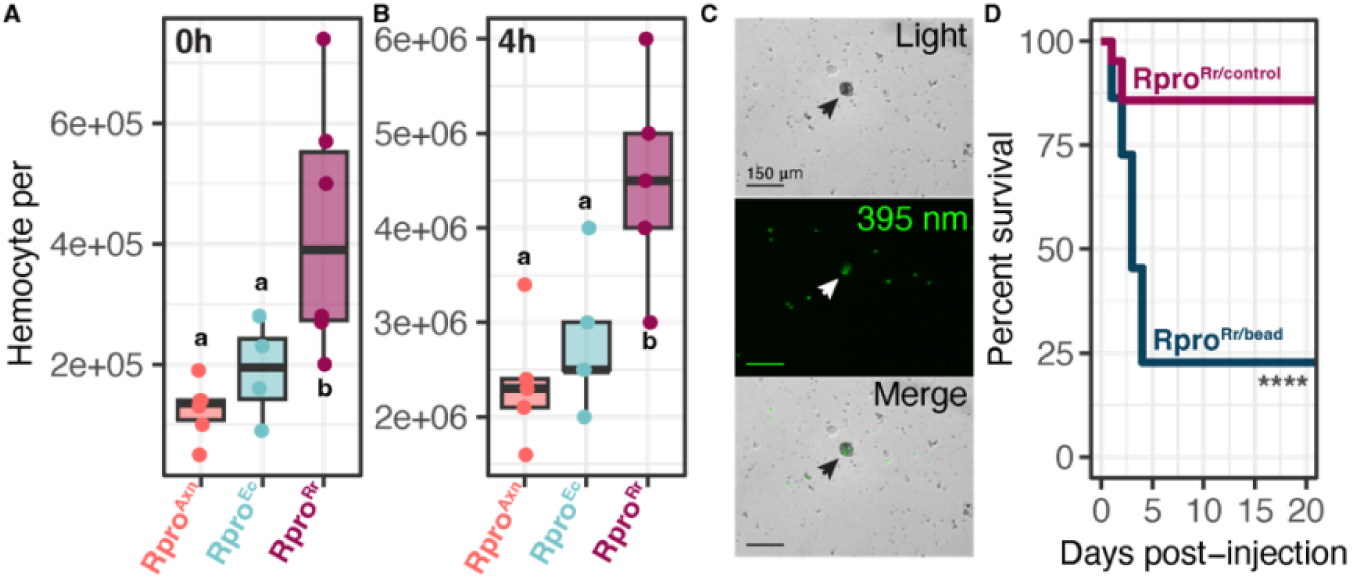
Rpro^Rr^ bugs have higher baseline and induced numbers of hemocytes relative to Rpro^Ec^ and Rpro^Axn^ bugs. **(A)** Hemocyte counts from hemolymph extracted at 0 h post-challenge with *E. coli*. There was no significant difference in the number of hemocytes between Rpro^Axn^ and Rpro^Ec^ but Rpro^Rr^ had significantly more than either (p = 0.0001, p = 0.02, Tukey’s HSD). **(B)** At 4h post challenge, all bugs had more hemocytes but Rpro^Rr^ still had significantly more than Rpro^Ec^ or Rpro^Axn^, which were not significantly different from one another (p = 0.0008, p = 0.001, p = 0.23 respectively, Tukey’s HSD). **(C)** Fluorescent latex beads are consumed by a hemocyte (arrow) in Rpro^Rr^. Top panel: light microscopy, middle panel: 395 nm, bottom panel: merge. **(D)** Hemocytes are essential for *R. rhodnii*-mediated immune effects. Inactivation of hemocytes by injection of latex beads dramatically reduced survival of Rpro^Rr^ bugs after challenge with *E. coli* (**** p < 0.0001, log-rank test).

The third pattern we observed was further evidence of crosstalk between the Toll and Imd immune pathways, consistent with the earlier experiments examining *dorsal* and *relish* (Fig. 3). Defensins, AMPs which act primarily against Gram-positive bacteria [46], were highly expressed in Rpro^Ec^ in response to challenge with Gram-negative bacteria (Fig. 5, *defensin 84, 82*). Likewise, *lysozyme42* expression was also induced by Gram-negative bacteria (Fig. 5) despite canonically being considered an AMP against Gram-positive bacteria [47]. A similar pattern was seen with expression of *prolixicin*, an ortholog of *Drosophila diptericin,* as it was induced in Rpro^Rr^ bugs after challenge with *M. luteus*. This is despite Diptericin having been shown to be active against Gram-negative bacteria [48]. Our data supports and expands on earlier work in the stink bug *Plautia stali* which demonstrated that expression of *relish* and several immune effectors can be induced by both *E. coli* and *M. luteus* in Hemiptera [45].

### *R. rhodnii* Influences the Cellular Immune System

Since the microbiome enhanced the humoral immune response and provided protection against bacterial infections, we wondered if gut bacteria-mediated immune priming extended to cellular immune responses. We first measured the number of circulating hemocytes in the hemolymph of Rpro^Rr^, Rpro^Ec^, and Rpro^Axn^ bugs. Fourth instar nymphs were immune challenged with *E. coli* and hemolymph was collected at 0 and 4 hours post-injection and a hemocytometer was used to count the number of hemocytes present. Immediately after injection Rpro^Rr^ bugs had a significantly higher number of circulating hemocytes compared to Rpro^Axn^ bugs (p = 0.0001, Fig 5A) and Rpro^Ec^ bugs (p = 0.02, Tukey’s HSD) and there was no significant difference in hemocyte counts between Rpro^Axn^ and Rpro^Ec^ (p = 0.11, Tukey’s HSD). These results demonstrate that Rpro^Rr^ bugs have a higher initial number of hemocytes than Rpro^Ec^ or Rpro^Axn^. Four hours after injection, all hemocyte titers had increased (Fig. 4B, p < 0.001 for all comparisons, Tukey’s HSD), but Rpro^Rr^ insects still had significantly higher hemocyte counts than both Rpro^Axn^ and Rpro^Ec^ bugs (p = 0.0008 and p = 0.001, respectively, Tukey’s HSD).

To investigate whether the differences in hemocyte counts among the treatments was a major factor in infection outcome, we suppressed hemocyte phagocytosis by pre-injecting fluorescent latex beads into the hemocoel of Rpro^Rr^ bugs [49]. The beads were visualized to be phagocytosed by the hemocytes *in vitro* (Fig. 5C). Twenty-four hours after bead injection, *E. coli* was injected into the hemolymph and survival was monitored as previously described. Insects with suppressed cellular immunity (Rpro^Rr/bead^) succumbed to bacterial infection faster than Rpro^Rr/control^ insects (p < 0.05, log-rank test, Fig. 5D), indicating that *R. rhodnii*-mediated cellular immunity plays an important role in immune priming, but that *E. coli* in the gut does not induce as strong of a cellular response.

### Melanization and Phenol Oxidase Activity are Dependent on the Presence of *R. rhodnii*

We observed throughout our experiments that Rpro^Axn^ individuals exhibited reduced wound healing. Rpro^Axn^ bugs did not develop a robust, dark melanization scar at injection sites but rather a light, thin scar, while Rpro^Rr^ developed a characteristic think, dark, scar (Fig. 6A). Interestingly we also saw a lack of dark, thick scar tissue in Rpro^Ec^ bugs. The dark scars following wounding is due to deposition of melanin produced by the action of phenol oxidases and other enzymes in the hemolymph which convert tyrosine to melanin [49]. The cascade leading to melanization can be triggered by both wounding and MAMPs [50]. Based on these observations and the role of melanization in insect immunity, we decided to further investigate differences in melanization by measuring phenol oxidase activity in Rpro^Rr^, Rpro^Ec^, and Rpro^Axn^ insects.

**Figure 6:**
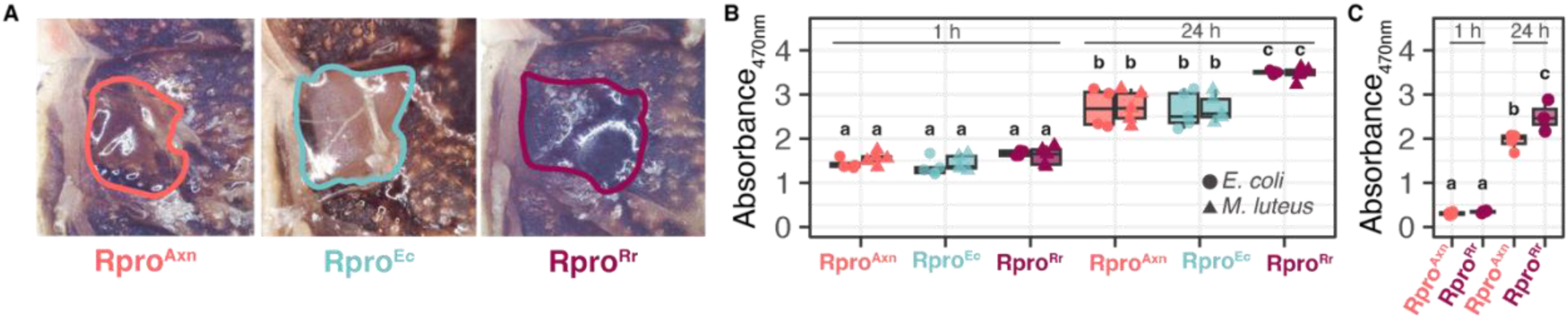
*R. rhodnii* enables successful melanization in *R. prolixus.* **(A)** Melanization of wounds is impaired in Rpro^Axn^ and Rpro^Ec^ relative to Rpro^Rr^. Wound area is outlined. **(B)** Hemolymph DOPA conversion assay of Rpro^Axn^, Rpro^Ec^, and Rpro^Rr^ performed at 1 or 24 h post-challenge with *E. coli* (circles) or *M. luteus* (triangles). At 0 h, there was no significant difference in conversion of DOPA to melanin between the bugs or bacterial challenges (F2,44 = 0.11 p = 0.74, ANOVA). At 24 h, there was a significant increase in DOPA conversion in all treatments relative to 0 h (p < 0.05, Tukey’s HSD). Rpro^Rr^ bugs had higher DOPA conversion than Rpro^Axn^ and Rpro^Ec^ bugs (p < 0.05, Tukey’s HSD), which were not significantly different from each other (p > 0.05, Tukey’s HSD). As with the 0 h time point, there was no significant effect of the bacterial species used in the challenge (F1,44 = 0.21, p = 0.648, ANOVA). **(C)** Wounding induces activation of the melanization response at 24h post-wounding. Rpro^Rr^ had higher DOPA conversion in response to wounding than Rpro^Axn^. Bars connected by different letters are significantly different (p < 0.05, Tukey’s HSD).

We measured phenol oxidase activity via a dihydroxyphenylalanine (DOPA) conversion assay [51] to assess how the presence of gut microbes influences melanization. DOPA is a precursor metabolite that is converted into melanin via the action of phenol oxidases. Hemolymph DOPA conversion was measured at 1 and 24 hours post injection with either *E. coli* or *M. luteus*. There was a highly significant effect of the gnotobiotic state (Fig. 6B, F_2,54_ = 23.8, p = 3.88 x 10^-8^, ANOVA) and time (F_1,54_ = 681, p < 2 x 10^-16^, ANOVA) on the amount of DOPA conversion. Surprisingly, there was not an effect of the microbe injected, as both *E. coli* and *M. luteus* injection triggered similar melanization among the different gnotobiotic states (Fig. 6B, F_3,54_ = 0.09, p = 0.96). There was no significant difference in DOPA conversion at 1h post-injection, but at 24 h post-injection there was a significant increase after injection of either *E. coli* or *M. luteus*. Compared to Rpro^Ec^ and Rpro^Axn^, Rpro^Rr^ had greater DOPA conversion regardless of challenge. We also investigated whether a stab wound alone was sufficient to trigger an increase in melanization. We stabbed Rpro^Axn^ and Rpro^Rr^ bugs with a sterile insulin syringe, then collected hemolymph at 1 and 24 h post-stab and measured DOPA conversion as before. We found that similar to injections with *E. coli* or *M. luteus*, stabbing with a sterile needle induced the ability to convert DOPA in both Rpro^Axn^ and Rpro^Rr^ hemolymph at 24 h (Fig. 6C, p < 0.001, Tukey’s HSD) but not at 1 h (p = 0.99, Tukey’s HSD), with a larger increase in conversion seen in the Rpro^Rr^ bugs (p = 0.014, Tukey’s HSD). Our DOPA conversion assays suggest that *R. rhodnii* is important for successful melanization in *R. prolixus*.

## Discussion

We demonstrate that the presence of gut microbes is integral to a functional immune system in *R. prolixus* and key to their ability to overcome infection with facultatively pathogenic microbes. We provide multiple lines of evidence that gut microbes in *R. prolixus* induce a stronger immune response, or “primes” the immune system by influencing humoral and cellular immunity, and that symbiotic *R. rhodnii* induce a stronger priming and protective effect than commensal *E. coli*. Bugs that harbored a microbe in their gut exhibited a strong humoral immune response by expressing significantly higher levels of antimicrobial peptides than microbe-free Rpro^Axn^ individuals, that corresponded to higher antimicrobial activity in their hemolymph, and subsequently an increased ability to reduce the number of infecting microbes compared to Rpro^Axn^ bugs. Our results suggest that different gut microbes prime the immune system in different ways and to different extents. *R. rhodnii* produces a stronger effect and modulates both cellular and humoral immunity while *E. coli* mainly stimulates humoral immune factors and does not have as strong of a priming effect.

Immune priming has been revealed to be a broadly important facet in insect immunity even though insects lack an adaptive immune system as seen in vertebrates. Despite the lack of antibody-mediated immune memory, in many insect species prior exposure to a nonlethal dose of a pathogen or pathogen derived material results in an elevated immune response which can be seen in the form of increased amount of AMPs and circulating hemocytes [52]. This elevated immune response results in the insect being resistant to a subsequent infection [18,53–56]. Immune priming has primarily been described in scenarios where an insect is exposed to an initial, non-lethal infection followed by subsequent exposure to the initial pathogen, known as homologous immune priming [57]. More recently, heterologous immune priming, in which one pathogen induces a protective immune response towards a different secondary pathogen has been observed [53]. Our results suggest that both the Gram-positive symbiotic *R. rhodnii* and Gram-negative commensal *E. coli* are capable of stimulating heterologous immune-priming against a different bacterial species, though to varying degrees.

Immune priming in insects has been thoroughly examined in several lineages including *Drosophila melanogaster* [58,59], *Aedes aegypti* [60], *Anopheles sp.* [61–63]*, Tenbrio* [64], *Gallaria mellonella* [65], *Tribolium castaneum* [66] and others. Inoculation of *D. melanogaster* with sub-lethal doses of *Streptococcus pneumoniae* led to life-long resistance to subsequent *S. pneumoniae* infection, which, similar to our observations required Toll activation and was mediated by hemocytes [64], however dead *S. pneumoniae* were sufficient to stimulate priming in *Drosophila* while dead *R. rhodnii* are not sufficient for *R. prolixus* priming. In *Drosophila* and *Aedes* mosquitos, *Wolbachia* has been demonstrated to upregulate immune-related genes of hosts which then confers protection against a variety of pathogens [67–71]. The mosquito gut symbiont *Asaia* has been shown to activate AMP production in the gut of *Anopheles stephensi* mosquitos [72]. We also observe an increase in expression of various immune genes when we alter the gut microbiome of kissing bug nymphs, though the identity and degree of expression varies based on the identity of the gut microbe.

The repeated evolution of symbiosis between insects and bacteria, in conjunction with diverse pathogenic threats, has resulted in a diverse array of mechanisms underpinning immune priming. In the bean bug, *Riptortus pedestris,* their gut symbiont *Caballeronia* strongly contributes to the activation ofs host immunity by influencing AMP activity and hemocyte number [65]. Other bacteria have been demonstrated to contribute to *R. pedestris* immune priming. A soil-derived *Burkholderia* colonizes *R. pedestris* and escapes the gut where it primes the immune system through influencing AMP expression and hemocyte number. This immune priming directly results in increased survival of *Burkholderia*-harboring insects compared to bugs who harbor *Caballeronia* alone [68]. We did not detect *R. rhodnii* outside of the *R. prolixus* gut, suggesting that a different mechanism is at work in kissing bugs.

Tsetse flies (*Glossina* sp.), which are also obligately exclusively hematophagous, harbor an intracellular symbiont, *Wigglesworthia glossinidia*. The symbiont is essential for proper immune function in *Glossina morsitans*, as removal of the symbiont leads to disruption of the immune response and melanization [17,73,74]. Like *R. prolixus*, absence of symbiotic bacteria leads to a reduction in AMP expression, loss of melanization response, and fewer circulating hemocytes. Interestingly, the melanization response appears to be linked to the activity of an odorant binding protein [74]. The loss of *Wigglesworthia* reduces expression of a peptidoglycan recognition protein, *pgrp-lb*, which in turn leads to activation of Imd signaling, while RNAi-mediated suppression of *pgrp-lb* in flies with their native *Wigglesworthia* also leads to Imd activation and AMP production, followed by a reduction in *Wigglesworthia* titers in the fly [75]. In *R. prolixus*, we observe higher expression of AMPs in Rpro^Rr^ bugs, and suppression of the Imd or Toll pathway leads to higher mortality after immune challenge, suggesting that while similarities exist between these two systems, the immune priming effect of *R. rhodnii* in *R. prolixus* differs in important ways from that of *Glossina* and *Wigglesworthia*.

Several mechanisms by which symbiotic bacteria protect their hosts from pathogens have been described. Competitive exclusion is one mechanism, as seen in honeybees where their native gut bacteria prevent opportunistic pathogens from colonizing the gut [76]. Symbionts can also directly interfere with pathogens by producing toxins that act against them [77]. We doubt this is a plausible mechanism in *R. prolixus*, as our experiments tested an immune response to bacteria directly injected into the hemocoel, bypassing the gut altogether and preventing direct contact between the microbes.

The effect of symbionts on immunity may be indirect, potentially mediated by nutritional factors. Rpro^Axn^ individuals lack symbiont-derived B vitamins, which are important for general homeostasis and may be essential for immune processes. *R. prolixus* relies on its microbiome to supplement B vitamins, key nutrients that are lacking in vertebrate blood [22,78]. Rpro^Axn^ individuals that were reared in axenic conditions suffer from increased development time, increased mortality, and failure to reach the reproductive adulthood stage [22,25,79]. The decrease in fitness seen in Rpro^Axn^ bugs potentially contributes in some way to their compromised immune function. Immune function has been linked to nutrition in many different insects, starving or rearing insects on nutritionally poor diets alters the humoral and cellular immunity leading to increased mortality after infection [80–82].

Diet has been shown to interact with the immune response in *R. prolixus* as well. Thirty days of starvation post-ecdysis resulted in increased mortality after infection due to changes in the cellular immune system, and similar results were found when bugs were fed an incomplete diet of plasma alone [80]. All bugs used in our experiments were two weeks post-molt to control for any effects starvation has on survival after infection. We attempted to disentangle the impacts of nutrition on survival by clearing Rpro^Rr^ with a blood meal containing antibiotics. These cleared bugs were confirmed to have no *R. rhodnii* present and were challenged with *E. coli*. Interestingly, these bugs suffered high mortality rates, though not as high as Rpro^axn^ bugs, suggesting that *R. rhodnii*’*s* influence on the immune system is not entirely mediated by nutritional factors. Conversely, axenic individuals that were fed a blood meal containing *R. rhodnii* at their 3^rd^ instar then challenged with *E. coli* shortly after had higher survival than Rpro^axn^ but significantly lower survival than our Rpro^Rr^ group. These results taken together suggest that nutritional supplementation via the microbiome may play some role in immunity, but likely other factors are contributing to this interaction. Direct experiments with B vitamin supplementation and B vitamin auxotrophic *R. rhodnii* will be necessary to fully resolve the role of symbiont-provisioned nutrients in *R. prolixus* immunity.

Another indirect mechanism of immune priming is the activation of humoral immune responses by gut bacteria. In the current study we demonstrated that the presence of the gut microbiome influenced these induced humoral immune responses. Naïve, unchallenged Rpro^Rr^ individuals had higher expression of both *relish* and *defensin82* compared to Rpro^axn^ individuals, suggesting that the presence of *R. rhodnii* is inducing an immune response. Interestingly, Rpro^Ec^ individuals had higher expression of all genes tested when compared to Rpro^axn^ bugs and higher expression than Rpro^Rr^ bugs for multiple genes tested. This suggests that different gut microbes activate the humoral immune system, and that some bacteria can induce humoral immunity more than the symbiont *R. rhodnii* alone. This does not translate to equivalent survival upon pathogen challenge, suggesting that other factors beyond expression of immune genes are playing key roles in symbiont-mediated immune priming.

The heterologous priming phenomenon seen in *R. prolixus* may be due to non-canonical activation or cross-talk between the Imd and Toll pathways. Immune signaling in insects was initially described in *Drosophila* as a linear response where different classes of pathogens triggered different immune pathways: Gram-negative bacteria activated the Imd pathway, Gram-positive bacteria and fungi activate the Toll pathway [83]. Yet as insect immune studies have moved beyond *Drosophila* and into other insects, it appears that crosstalk between immune pathways may be more common than previously thought. Work in the hemipteran stink bug, *Plautia stali*, revealed that not only are both Imd and Toll pathways present but there is a blurred functional differentiation, as immune challenge with Gram-negative or Gram-positive bacteria elicited expression of immune effector genes of both pathways [82]. We see similar patterns in our expression data as immune challenge with Gram-positive and Gram-negative trigger expression of both *dorsal* and *relish* immune transcription factors belonging to the Toll and Imd pathway respectively. Knockdown of either *dorsal* or *relish* also result in increased mortality after challenge with Gram-negative *E. coli,* and AMPs thought to be active against either Gram-positive or Gram-negative bacteria are expressed in response to both Gram-negative and positive bacteria. Crosstalk has been seen in other insects, including the beetle *Tenebrio molitor* [83], and other species of kissing bugs. In *Triatoma pallidipennis,* silencing of Toll pathway genes led to increased mortality when bugs were challenged with a Gram-negative bacteria [84]. Interestingly, silencing of *relish* in *T. pallidipennis* did not result in increased mortality. Even in *Drosophila*, gut bacteria can elicit immune priming against heterologous pathogens including fungi [84] and viruses [85].

Our results contribute to a growing understanding that Hemipteran immunity seems to function differently than many holometabolous model organisms such as *Drosophila*. Genomic data has revealed that many hemipteran genomes lack elements of the canonical insect immune pathways, including aphids [86], bedbugs [87], and scale insects [88]. The genome sequence of *R. prolixus* was initially thought to be missing key components of the Imd pathway [89], though subsequent analysis has revealed that despite lack of the genes *imd* and *kenny*, *R. prolixus* does indeed possess a functioning Imd pathway [90]. Our results support these earlier findings, as both *R. rhodnii* and *E. coli* stimulate expression of relish and Imd-associated AMPs, while inactivation of Imd signaling through RNAi silencing of relish lead to increased mortality of *R. prolixus.* The absence of *imd* and *kenny* suggests that Imd immune signaling functions differently in *R. prolixus* and more research is needed to further understand immune signaling in *R. prolixus*.

Most insects possess an acellular chitinous and proteinaceous peritrophic matrix that lines the midgut epithelium and is responsible for protecting the midgut cells from direct contact with gut microbes. This protective barrier modulates immune activation by the gut microbiome [91]. Hemipterans, including kissing bugs, do not have a peritrophic matrix (PM), but rather a lipid-based structure called the perimicrovillar membrane (PMM). While the PMM forms a barrier between the gut lumen and its resident microbiota, it is possible that the PMM is not as significant a barrier to microbes or MAMPs, and may permit direct contact of gut microbes with the gut epithelia. Such direct contact could potentially activate host immune responses to a greater extent than if bacteria did not contact the epithelial cells directly [92]. Additional studies are necessary to understand the extent to which gut bacteria or MAMPs in *R. prolixus* encounter the epithelium, and how this influences immune responses.

Kissing bugs are the vectors of *Trypanosoma cruzi*, the causative agent of Chagas disease. *T. cruzi* is a stercorarian parasite, residing exclusively in the gut of its triatomine host during the insect phase of its development. As a result, it may be directly or indirectly influenced by the presence of bacteria. The activation of the immune system by gut bacteria may have consequences for *T. cruzi* persistence and transmission, as has been seen in *Anopheles* mosquitoes and their gut bacteria [93], where presence of gut bacteria induces a strong immune priming effect via hemocyte differentiation following ookinete escape from the midgut, and subsequently reduces the survival of *Plasmodium* in the mosquito.

In kissing bugs, the microbiome has been implicated in both interactions with the host immune system as well as antagonistic interactions with *T. cruzi*[94–99]. Batista et al found similar results to ours, with restoration of *R. rhodnii* to antibiotic-cleared *R. prolixus* leading to increased expression of AMPs. Likewise, inactivation of Imd signaling with a pharmacological agent, IMD-0354, lead to increased mortality in *R. prolixus* due to infection [95]. These experiments were focused on the outcomes of bacterial infection in the gut, while ours examined infection of the hemocoel. Together they suggest that the immune priming effect of symbiotic bacteria also likely occurs in the gut. Our results support and expand upon these previous studies and provide a framework for further understanding how gut microbiomes influence kissing bug immunity and *T. cruzi* transmission.

Our study provides evidence that the gut microbiome in kissing bugs plays an essential role in activating the host immune system against pathogens in the hemocoel. The nature of the immune priming appears to vary based on the identity of the gut microbe in question, as symbiotic microbes provide a stronger protective effect than non-symbiotic commensals. Surprisingly, both commensal and symbiotic bacteria were able to activate both the Toll and Imd pathways, revealing that our understanding of insect immunity in kissing bugs and possibly other hemipterans, largely based on studies in *Drosophila* and other holometabolous insects, is not complete.

## Materials and Methods

### Insect Maintenance

*Rhodnius prolixus* were obtained from the lab of Dr. Ellen Dotson at the Centers for Disease Control and Prevention through BEI Resources. Insects were reared at 28 °C with a photoperiod of 12 h of light and 12 h of dark and 80% relative humidity. General colony insects were kept in 1 L Nalgene containers and regularly fed defibrinated rabbit blood (Hemostat Laboratories, Dixon, CA) inoculated with *R. rhodnii* bacteria in the exponential phase of growth through an artificial membrane feeder.

### Generation of Axenic and Gnotobiotic Nymphs

*R. prolixus* eggs were collected 7 days after being laid then placed in a sterile cell collection basket and washed with 70% ethanol for 5 minutes followed by 3 minutes in 10% povidone-iodine solution, then another 5-minute wash in 70% ethanol, followed by three rinses in autoclaved deionized water. Sterilized eggs were then transferred to autoclaved glass containers enclosed in sterile Nalgene containers with gas-exchange tape covering an air hole. Sterility was validated by screening total genomic DNA from insects with PCR to amplify a 16S rDNA gene with the universal primers 27F and 1492R (supplementary table S1). No bands were observed in axenic nymphs. Gnotobiotic nymphs were generated by feeding axenic first instar nymphs a blood meal inoculated with 10^6^ CFU/mL of *R. rhodnii or E. coli*. Nymphs were fed at every instar approximately 2 weeks after molting. Gnotobiotic states were confirmed through qPCR on DNA extracted from whole bodies of nymphs using primers specific to the *gyrB* sequence of each bacteria (supplementary table S1).

### Bacterial Strains

*Rhodococcus rhodnii* (NRRL B-16535) was obtained from ATCC and grown at 28 °C in liquid Luria-Broth (LB). *Escherichia coli* MG1655 was a gift of Eric Stabb and was grown in liquid LB at 37 °C. *Micrococcus luteus* NCTC 2665 bacteria was a gift of Michael Strand and was grown in liquid LB at 37 °C. Bacterial titers were determined by measuring the OD_600_ of cultures on a Beckman Coulter DU640 spectrophotometer and then plating out serial dilutions of culture on LB agar plates to correlate OD_600_ with Colony Forming Units (CFU) counts.

### Bacterial Immune Challenge

Bacterial immune challenge in kissing bugs was performed on either Rpro^axn^, Rpro^Ec^, or Rpro^Rr^ 4^th^ instar nymphs that were two weeks post molt. Bacteria were injected intrathoracically with 2 μl of 10^8^ CFU/ml of an overnight culture. Nymphs were challenged with either *R. rhodnii*, *M. luteus*, *E. coli*, or sterile *Aedes* saline. Bacteria were collected by centrifugation at 8,000 x g for 5 minutes and resuspended in sterile *Aedes* saline. A group of nymphs received a stab wound without injection and a group of nymphs were unaltered and left as a naïve treatment group. Twenty-four hours after injection, whole guts and fat body were dissected out in sterile PBS and stored at-80 °C.

### Real-Time Quantitative PCR (qPCR)

For analysis of gene expression, axenic and gnotobiotic individuals’ total RNA was isolated from homogenized tissues using the Direct-zol 96 RNA MagBead Kit (Zymo Research) and KingFisher Apex extraction system. Total RNA was subject to DNAse treatment with the Turbo DNA-free kit (Thermo Fisher Scientific) according to manufacturer’s instructions. Purified, DNased RNA quantification and purity was validated using a NanoDrop spectrophotometer to measure absorbance ratios. One hundred nanograms of RNA was reverse transcribed using iScript Reverse Transcription Supermix (Bio-Rad). qPCR was performed on synthesized cDNA in quadruplicate using the QuantiNova SYBR Green master mix (Qiagen) in a total volume of 20 µl with 0.5 mM of each primer on a Roche LightCycler96 system or a Qiagen Rotor Gene system. For each gene tested, 4 technical replicates and 5 biological replicates were performed. Absolute quantification of genes was performed as previously described [94] using standard curves of pSCA plasmids containing qPCR products. All qPCR primer pairs had an efficiency of > 0.85.

### Survival Analysis

*E. coli*, *R. rhodnii*, and *M. luteus* were grown overnight at 37 °C in LB broth to OD_600_ = 1. Cells were centrifuged at 6,000 rpm and resuspended in sterile *Aedes* saline to a concentration of 5 x 10^7^ CFU/ml. Fourth instar Rpro^axn^, Rpro^Rr^, or Rpro^Ec^ nymphs 2 weeks post-molt were injected with 2 µl (10^6^ CFU) of either *E. coli*, *R. rhodnii*, *M. luteus*, or sterile saline with a sterile Hamilton syringe using a Micro4 syringe pump controller (World Precision Instruments). Nymphs were placed individually into wells of a sterile 24-well polystyrene cell culture plates for observation and mortality was observed daily for 21 days post-injection.

### Bacterial Clearance in Hemolymph

Bacterial abundance in hemolymph was measured by collecting hemolymph from 4^th^ instar nymphs after challenge with kanamycin resistant *E. coli* as described above. All legs were removed with forceps, then an individual was placed inside a sterilized, filtered p1000 pipette tip inserted into a 2 mL microcentrifuge tube, which was then centrifuged at 2000 RCF for 10 min at 15 °C, resulting in the collection of 2-3 µl of hemolymph from an individual. Hemolymph of 4 individuals per treatment was pooled, diluted in 20 µl of sterile PBS, and spread on LB agar plates with 50 µg/ml of kanamycin sulfate, then incubated for 24 h at 37°C and the number of CFUs were counted. For each treatment, three biological replicates were performed.

### Quantification of Antimicrobial Hemolymph Activity

Antimicrobial activity of kissing bug hemolymph was measured using a zone of inhibition assay as described by [42,43]. A culture of *M. luteus* was grown overnight in LB at 37 °C, then 1 mL of culture was added to 10 mL of sterile, cooled liquid LB agar. The *M. luteus*-LB agar solution was mixed and poured into Petri plates. After solidifying, 1 mm diameter holes were created in the agar using a sterile glass Pasteur pipette. Hemolymph was collected from individual nymphs as described above, and 1 µl of hemolymph was placed in each hole and the plates were incubated overnight at 37 °C. The diameters of the individual zones of bacterial growth inhibited were measured using an ocular micrometer on a stereo dissecting microscope. Two independent trials were conducted, and each trial consisted of 10 bugs per treatment group.

### Antibiotic Clearing/Recolonization of *R. rhodnii*

To investigate the impacts on survival of microbiome recolonization, we inoculated 3^rd^ instar Rpro^axn^ bugs with *R. rhodnii* (Rpro^Axn + Rr^) through a blood meal as previously described. Bugs were allowed to develop to the 4^th^ instar when the presence of *R. rhodnii* was confirmed in a subset of bugs via qPCR (Fig. S4). Two weeks after molting bugs were immune challenged with *E. coli* and survival was monitored as previously described. To determine the effects of removal of *R. rhodnii* on host immune function, 3^rd^ instar Rpro^Rr^ were fed a bloodmeal containing either 150 µg/ml of kanamycin or a bloodmeal containing 150 µg/ml of kanamycin along with kanamycin resistant *R. rhodnii* (Rpro^Rr^ + Kan and Rpro^Rr-KanR^ + Kan). Bugs were confirmed to be removed of *R. rhodnii* or confirmed to still harbor *R. rhodnii* via qPCR as described above. All bugs were then allowed to molt to the 4^th^ instar then immune challenged as previously described and survival was monitored.

### RNAi-Mediated Immune Suppression

PCR Primers containing the minimal T7 promoter sequence were designed to amplify 400-500bp of *relish, dorsal, or egfp* (Table S1). Total RNA was extracted from the fat bodies of 4^th^ instar nymphs, DNased, and reverse transcribed as described above. The PCR product was subsequently cloned to the pSCA vector using the Strataclone PCR cloning kit (Agilent). Target DNA was amplified by PCR from isolated plasmid DNA. dsRNA was synthesized using the MEGAscript RNAi kit (ThermoFisher Scientific) according to the manufacturer’s instructions. Synthesized dsRNA was precipitated with sodium acetate and ethanol, then resuspended to 2 μg/μl in *Aedes* saline. Fourth instar nymphs were injected with 1 μl of dsRNA, then allowed to recover for 1 week before immune challenge as described previously. Knockdown of *relish* or *dorsal* was confirmed by qPCR on cDNA extracted from treated nymphs as described previously.

### Hemocyte Quantification and Inactivation

Hemolymph was collected from 4^th^ instar nymphs that were 2 weeks post-molt. Hemolymph was collected via perfusion of 100 µL of cold *Aedes* saline injected through the abdomen via insulin syringe. The samples were stored on ice until hemocyte numbers were counted using a Neubauer hemocytometer and an inverted stereo micrscope. Two counts per insect were conducted and the average of the two counts was used for each of 5 individuals per treatment.

To reduce hemocyte activity, fluorescent latex microbeads (Polyscience, Fluoresbrite Microspheres 2.00 µm) were diluted to 10^8^ beads/μl in *Aedes* saline and injected into the hemolymph of 4^th^ instar nymphs that were two weeks post-molt. Beads were confirmed to be engulfed by host hemocytes by observation with an epifluorescence microscope (Leica). Four hours after injection with beads, nymphs were challenged with injection of either *E. coli* or sterile *Aedes* saline then monitored for survival as previously described.

### DOPA Conversion Assay

Hemolymph was collected as described above from Rpro^axn^, Rpro^Ec^, or Rpro^Rr^ injected with either *E. coli* or *M. luteus*. DOPA conversion was measured as described in [100]. Briefly, 100 µl of perfused hemolymph was suspended into 100 µl of PBS containing 4 mg/ml DOPA, and added to the wells of a sterile 96 well plate. The plate was then incubated at 28 °C for 1 h in a humidified chamber and absorbance was read at 470 nm on a µQuant plate reader (BioTek). Four to six bugs per treatment were tested.

## Statistical Analyses

Statistical analysis of insect survival was determined using a Cox-proportional hazards model followed by a log-rank test for pairwise comparisons via R package *survminer* and *survival*. Bacterial clearance was analyzed using a Wilcoxon rank-sum test with a Benjamini-Hochberg correction for multiple comparisons. Statistical analysis of gene expression and hemocyte counts was performed using an aligned-rank transformed ANOVA test followed by a Tukey post-hoc test using the R package *ARTool* [100]. DOPA conversion assay data was analyzed via ANOVA and Tukey post-hoc tests. Data files and R scripts used in the study are found in the supplemental online data.

## Acknowledgements

We would like to thank Logan Blankenship, Bradley Mackett, Betsy Jackson, and Cassandra Armstrong for their assistance in maintaining our colony of *R. prolixus*. Dr. Michael Strand provided guidance on hemocyte quantification and inactivation experiments. Loretta Mugo and Sandra Mendiola provided helpful feedback on the manuscript.

---

**Figure S1:**
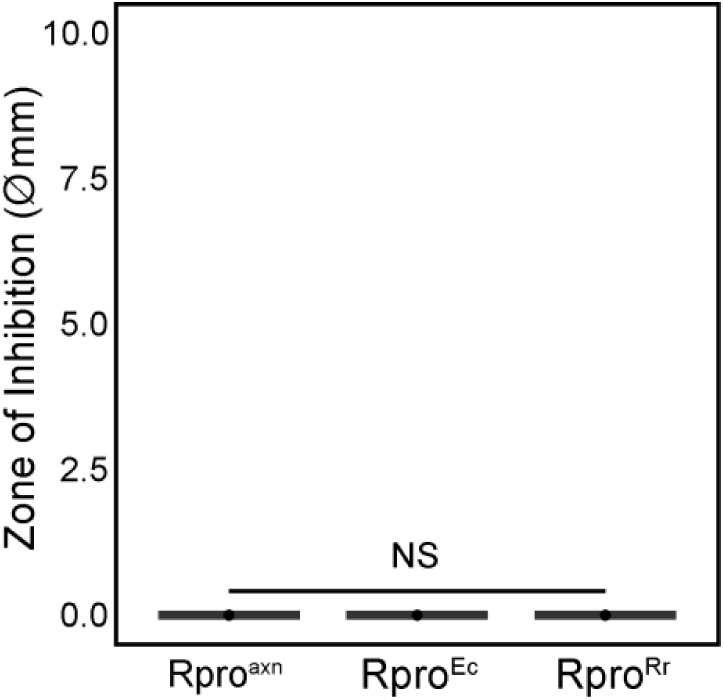
Zone of inhibition assay from naive insects. No inhibition of growth was observed in any gnotobiotic background.

**Figure S2:**
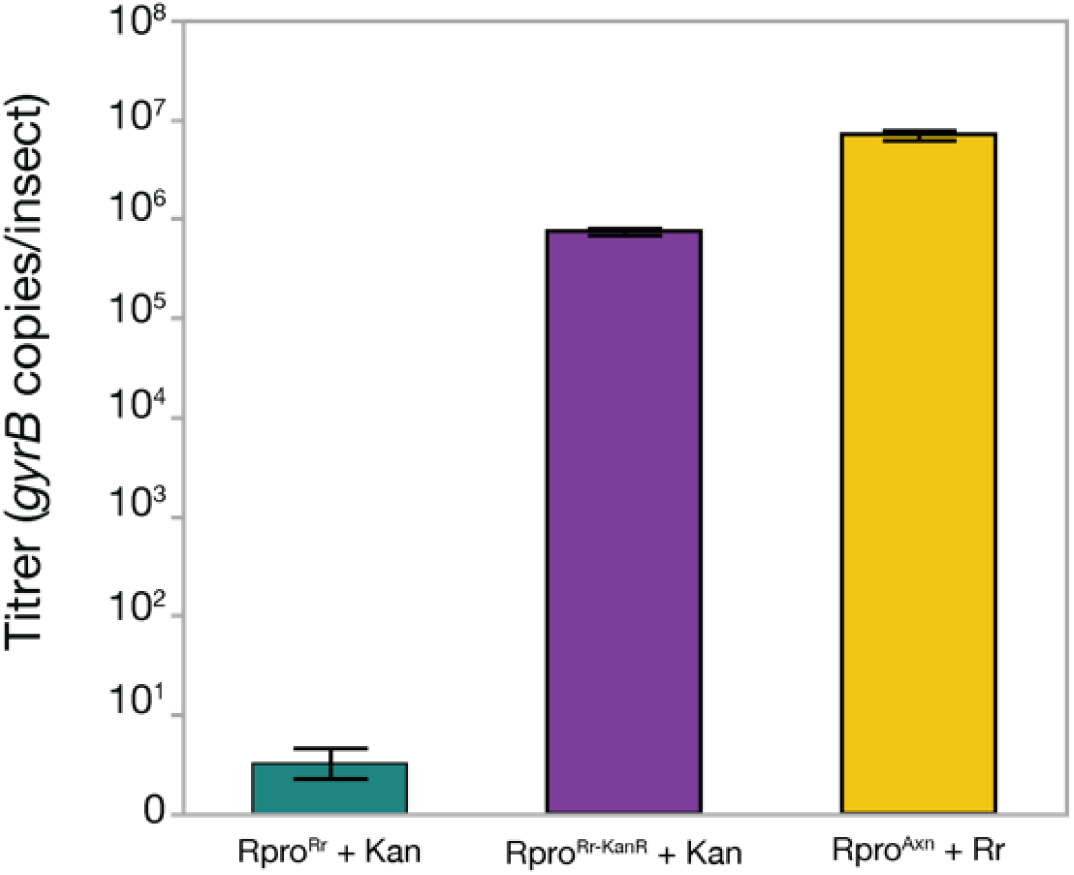
R *rhodnii* titer in Rpro^Rr^ and Rpro^Rr KanR^ insects treated with kanamycin (Rpro^Rr^ + Kan) and Rpro^Axn^ fed *R. rhodnii.* Titer was determined using qPCR to measure the number of *gyrB* copies per insect using extractions of whole-body DNA.

**Figure S3:**
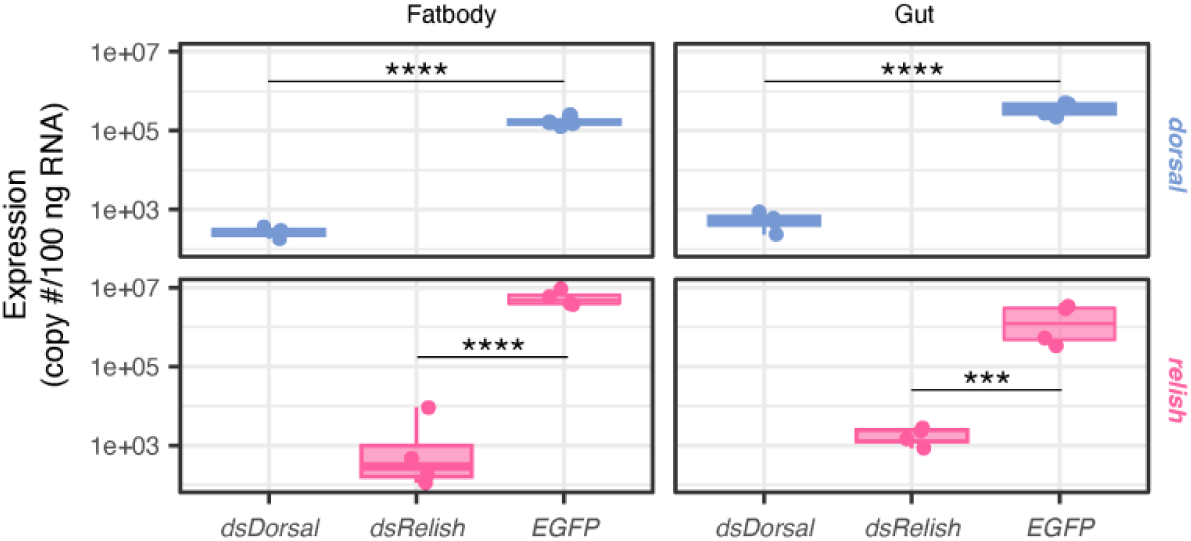
qPCR confirmation of dsRNA knockdown of *relish* and *dorsal*. RNA was extracted from either gut or fatbody. Asterisks indicate signficiant differences **** p < 0.001, *** p < 0.001, independent contrast test.

**Table SI:**
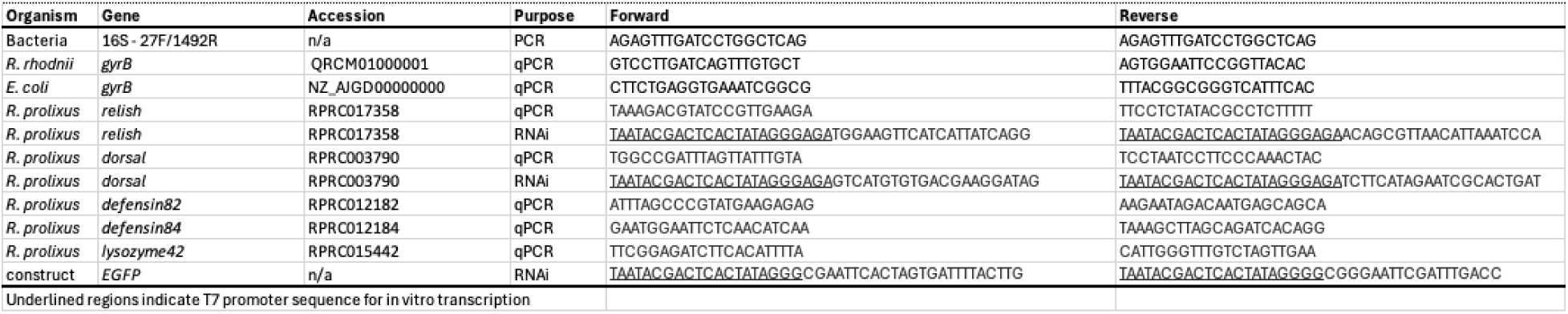
Primer sequences used in this study.

**Table S2:** Statistical details for gene expression experiments.

Underlined regions indicate T7 promoter sequence for in vitro transcription
| Table S2: Statistical details for gene expression experiments |  |  |  |  |  |
| --- | --- | --- | --- | --- | --- |
| Gene = dorsal | Df | Df.res | F | value | Pr(>F) |
| treatment | 2 | 26 | 29.802 | 1.87E-07 | *** |
| bug | 2 | 26 | 17.932 | 1.28E-05 | *** |
| treatment:bug | 4 | 26 | 60.4 | 8.76E-13 | *** |
| Gene = relish |  |  |  |  |  |
| treatment | 2 | 27 | 17.6574 | 1.25E-05 | *** |
| bug | 2 | 27 | 9.0321 | 0.0009924 | *** |
| treatment:bug | 4 | 27 | 8.7543 | 0.00011486 | *** |
| Gene = prolixicin |  |  |  |  |  |
| treatment | 2 | 27 | 49.85 | 8.63E-10 | *** |
| bug | 2 | 27 | 39.08 | 1.07E-08 | *** |
| treatment:bug | 4 | 27 | 47.585 | 7.59E-12 | *** |
| Gene = defensin84 |  |  |  |  |  |
| treatment | 2 | 27 | 11.536 | 0.00023925 | *** |
| bug | 2 | 27 | 10.728 | 0.00037261 | *** |
| treatment:bug | 4 | 27 | 33.441 | 4.24E-10 | *** |
| Gene = defensin82 |  |  |  |  |  |
| treatment | 2 | 25 | 11.755 | 0.00025202 | *** |
| bug | 2 | 25 | 19.18 | 8.94E-06 | *** |
| treatment:bug | 4 | 25 | 10.829 | 3.11E-05 | *** |
| Gene = lysozyme42 |  |  |  |  |  |
| treatment | 2 | 26 | 111.709 | 1.72E-13 | *** |
| bug | 2 | 26 | 38.496 | 1.69E-08 | *** |
| treatment:bug | 4 | 26 | 225.482 | < 2.22 E-16 | *** |

